# Chronic intestinal immune activation reveals separable impacts of inflammation and barrier loss on hallmarks of ageing

**DOI:** 10.1101/2025.08.11.669650

**Authors:** Jeanette Alcaraz, Charlotte Keyse, Charles Hall, David W. Walker, David P. Doupé, Rebecca. I. Clark

## Abstract

Inflammaging is considered a driver of age-associated pathology across tissues. Similarly, intestinal permeability is a feature of ageing and underlies a range of inflammatory and age-related diseases. Increased intestinal permeability has been described as both a cause and a consequence of inflammation. Both intestinal permeability and inflammation are closely associated with microbial dysbiosis, epithelial dysplasia and mortality but dissecting the complex interplay between these phenotypes remains challenging.

Here we genetically induce intestinal immune activation in *Drosophila* and stratify animals by their intestinal barrier status using the Smurf assay. We demonstrate that intestinal immune activation and barrier failure have distinct impacts on the microbiota. Further, intestinal immune activation drives intestinal barrier failure and mortality even in the absence of the microbiota. Importantly, immune-induced intestinal barrier failure takes time to develop and is closely associated with the onset of mortality.

Our work adds to building evidence that the impact of intestinal permeability on the microbiota and on animal health needs to be considered independently of its relationship with inflammation.

## Introduction

Chronic low-grade inflammation, or “inflammaging,” has been proposed as a unifying mechanism connecting age-associated pathologies across tissues (1). Elevated intestinal permeability has also been observed in a range of inflammatory and age-related diseases (2,3) and is increasingly recognized as a physiological feature of ageing. Intestinal permeability has been linked to systemic inflammation, microbial dysbiosis, and mortality across species (4–7). However, the directionality of the association between intestinal permeability and inflammation, and the role of the intestinal microbiota in this relationship, remains unclear. Specifically, it is not well established whether intestinal permeability initiates inflammation and pathology or arises as a secondary consequence (3,8).

It is well established that inflammation, via cytokines such as TNF-α and IFN-γ, can disrupt epithelial tight junctions and promote barrier leakiness (8,9). However, several longitudinal and mechanistic studies suggest that increased intestinal permeability may precede and contribute to the development of inflammatory disease. In human cohorts, elevated intestinal permeability has predicted subsequent development of Crohn’s disease (10,11). Similarly, age-associated increases in intestinal permeability and tight junction remodeling have been documented in non-human primates (12), indicating that epithelial barrier deterioration may occur independently of systemic inflammation. Evidence from multiple species supports that disruptions in junctional integrity and altered barrier function can be early features in both disease progression and ageing processes (13).

This bidirectional relationship between inflammation and intestinal permeability is further complicated by a close association with microbial dysbiosis and epithelial dysplasia. In model organisms these phenotypes are often co-regulated and experimentally inseparable, whereby interventions targeting one phenotype frequently impact the others (6,14–19). Intestinal permeability, age-related inflammation and increased microbial loads have been documented in aged flies, mice and monkeys (6,7,14). Germ free flies and mice show reduced age-related inflammation and improved barrier function (6,14) suggesting that the aged microbiota may drive both inflammation and intestinal permeability. However, studies in *Drosophila* have also demonstrated a role for innate immune activation in driving microbial dysbiosis. Immune signalling is normally tightly controlled by multiple negative regulators. A loss of negative regulation of immune signaling results in chronic immune activation, microbial dysbiosis and shortened lifespan (16). Conversely, restoration of immune suppression can prevent age-related microbial overgrowth and improve lifespan (17,18).

Dissecting these complex interrelationships requires experimental approaches that allow the separation of the effects of immune activation, epithelial dysplasia, intestinal permeability and microbial dysbiosis. Such approaches are essential if we are to move toward a causal understanding of the role of these phenotypes in age-related decline.

In *Drosophila*, the “Smurf” assay offers a well-established non-invasive method to stratify animals based on intestinal permeability. However, to date studies focused on the role of immune activation in driving microbial dysbiosis, epithelial dysplasia and mortality that measure or account for intestinal permeability are lacking. In this study, we use a *Drosophila* model of intestinal innate immune activation to investigate the relationship between immune activation, intestinal barrier dysfunction and associated phenotypes. We stratify animals by their intestinal barrier status allowing us to clarify the relationships between innate immune signaling, intestinal permeability, microbial dysbiosis, epithelial dysplasia and mortality.

## Results

### Intestinal overexpression of PGRP-Lc drives intestinal barrier loss and mortality

We previously demonstrated that overexpression of *PGRP-Lc* with the drug-inducible geneswitch driver 5966 drives mortality and loss of intestinal barrier function ((14), Figure S1A & B). PGRP-Lc is a receptor for the Immune Deficiency (IMD) pathway and its overexpression drives immune activation. 5966 drives *UAS-PGRP-Lc* expression in the intestine and in other tissues (Figure S1D & E) in adult females.

This results in IMD activation in multiple tissues, measured here with qPCR of the antimicrobial peptide Diptericin (Figure S1F & G). To further assess the impact of *PGRP-Lc* overexpression in different tissues we compared the ubiquitous geneswitch driver DaGS, the fat body and gut geneswitch driver s106, and the gut specific geneswitch driver TiGS. We found that *PGRP-Lc* overexpression drove early mortality and loss of intestinal barrier function in each case (Figure 1A-F, Fig S1C). Given that gut specific overexpression showed similar mortality to ubiquitous overexpression we conclude that *PGRP-Lc* overexpression in the intestine is sufficient to drive these phenotypes.

**Figure 1:**
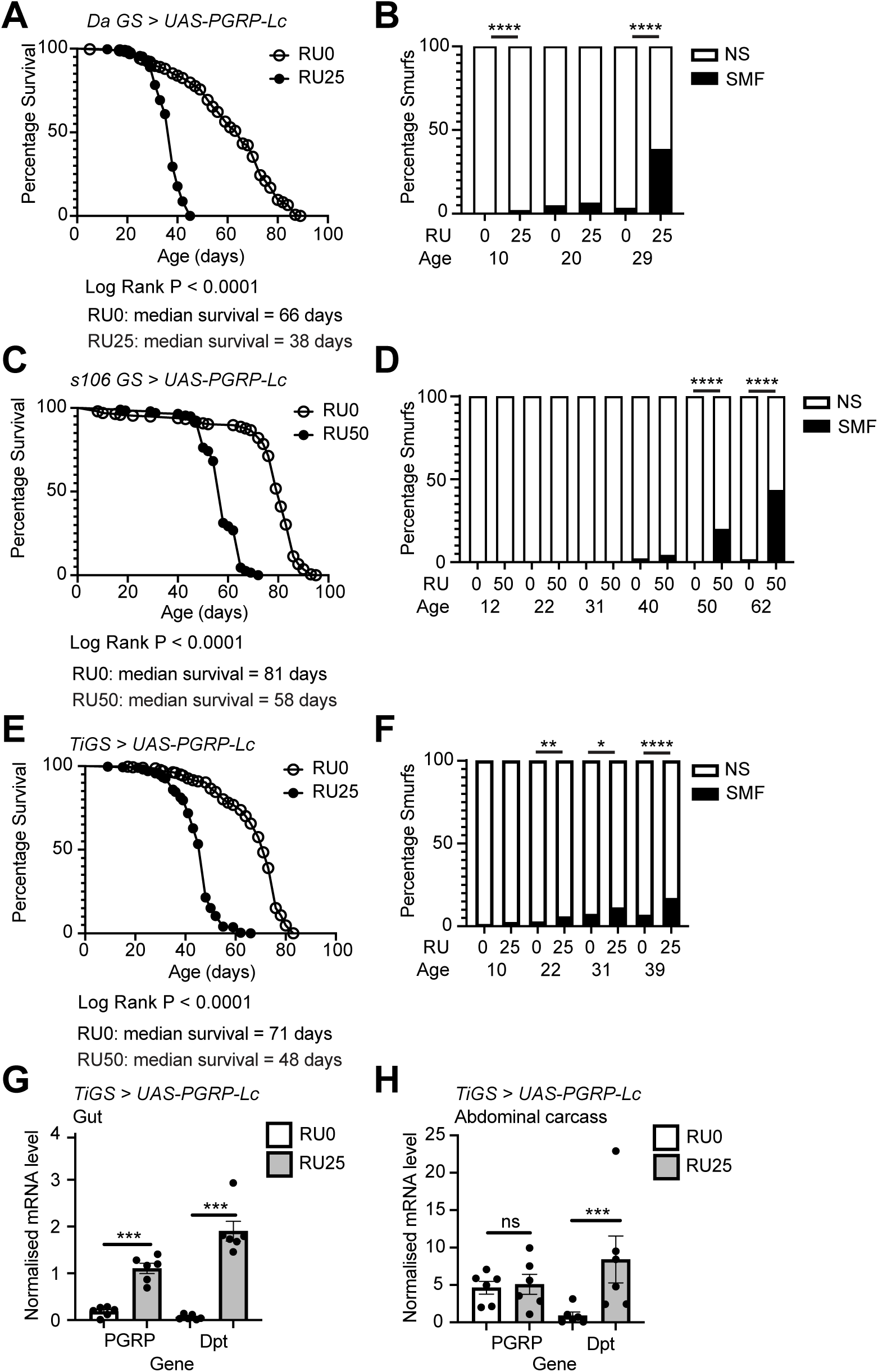
Intestinal *PGRP-Lc* overexpression drives early mortality and intestinal barrier loss. (A-F) Lifespan curves (A, C and E) and Smurf proportions (B, D and F) for *DaGS > UAS-PGRP-Lc* (A and B), *S106 GS>UAS-PGRP-Lc* ((C and D) and TiGS>UAS-PGRP-Lc (E and F) female flies drug induced (RU25, RU50) from early adulthood and uninduced controls (RU0). n > 200 flies/condition, NS = non-Smurf, SMF = Smurf. Log rank test was used for survival data and binomial test for Smurf proportions. (G-H) Normalised mRNA level for *PGRP-Lc* and *Diptericin (Dpt)* in dissected whole gut (G) and abdominal carcass samples. n = 6 samples, 5 guts/sample and 5 abdomens (gut removed)/sample. Bar graphs show mean ± SEM. Two-way Anova with Tukey’s multiple comparisons. *p < 0.05, **p < 0.01, ***p < 0.001, ****p < 0.0001.

However, while *PGRP-Lc* overexpression was restricted to the gut in *TiGS>UAS-PGRP-Lc* flies (Figure 1G & H), Diptericin expression was nevertheless strongly induced outside the gut (Figure 1G & H). Previous work has shown communication between local and systemic immune responses (20). This combined with our findings suggests that immune activation in the gut may always induce a systemic immune response. This would mean that it is not possible to induce immune activation only in intestinal tissue. It also means that doing so would not reflect normal physiology.

Given this it is important to note that we cannot distinguish the roles of intestinal and systemic immune activation in driving intestinal barrier loss and subsequent mortality. We have used both the 5966 and TiGS drivers to overexpress PGRP-Lc throughout this study and hereafter both are referred to as PGRP-Lc overexpression in the text. Which driver has been used is clearly stated on the figures and in the figure legends. Both drivers result in immune activation both in the intestine and systemically

### The impact of intestinal overexpression of PGRP-Lc on the microbiota is species/strain specific

Previous work has shown that Drosophila carry an increased internal microbial load with age and following intestinal barrier loss (4,14,15). Therefore, we assessed the impact of intestinal immune activation on microbial load. We found that overexpression of *PGRP-Lc* resulted in elevated internal microbial loads in adult female flies carrying their native microbiota (Figure 2A & B). Importantly, increased microbial load was observed in flies overexpressing *PGRP-Lc* that had not yet lost intestinal barrier function.

**Figure 2:**
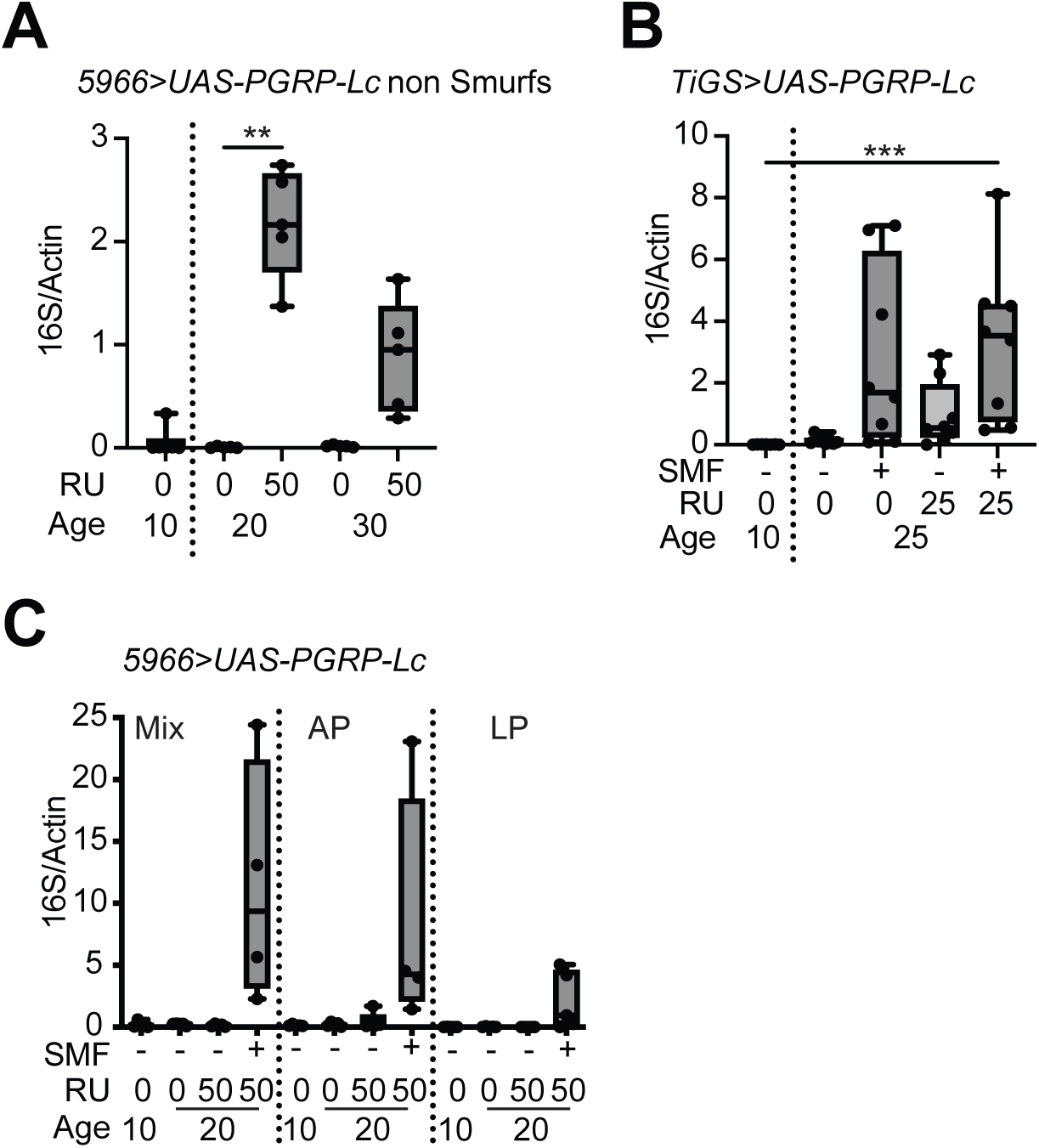
Intestinal overexpression of *PGRP-Lc* drives microbial growth in a species/strain specific manner. (A-C) Internal bacterial levels assayed by qPCR of the 16S rRNA gene in surface sterilised females flies drug induced (RU50) from early adulthood and uninduced controls (RU0). n = 6 samples with 5 flies/sample. (A) *5966>UAS-PGRP-Lc* non-Smurf flies, (B) *TIGS>UAS-PGRP-Lc* non-Smurf and Smurf flies and (C) *5966>UAS-PGRP-Lc* flies monoassociated with *Acetobacter pasteurianus* (AP), *Lactiplantibacillus plantarum* (LP) or associated with a mix of both species. Boxplots show display the 25-75^th^ percentiles, with the horizontal bar at the median, and whiskers extending from the minimum to maximum points. Two-way Anova with Tukey’s multiple comparisons. **p < 0.01, ***p < 0.001.

To clarify the impact of *PGRP-Lc* overexpression on different microbial groups we used flies monoassociated with two bacterial species commonly found in the *Drosophila* intestine, *Acetobacer pasteurianus* and *Lactiplantibacillus plantarum*, or associated with a mix of both species. Interestingly, we found that *PGRP-Lc* overexpression did not drive overgrowth of either of these bacterial species in flies that had not lost intestinal barrier function (Figure 2C). As expected from previously published work (4,14,15) flies that had lost intestinal barrier function showed elevated internal bacterial loads regardless of which bacteria were present and *PGRP-Lc* expression status (Figure 2B & C). We conclude that the impact of *PGRP-Lc*, i.e. IMD activation, on microbial growth is species or strain specific. This distinguishes the impact of immune activation on the microbial population from that of intestinal barrier loss, which creates conditions that enable overgrowth of every microbial species and population so far tested.

### Intestinal overexpression of PGRP-Lc drives intestinal barrier loss and mortality independent of the microbiota

To test the impact of microbial overgrowth on the reduced lifespan and early intestinal barrier loss that results from *PGRP-Lc* overexpression, we assessed these phenotypes in axenic, or germ-free, flies. Axenic flies overexpressing PGRP-Lc showed increased mortality and loss of intestinal barrier function like that of their conventionally reared counterparts (Figure 3A & B). While axenic flies overexpressing PGRP-Lc showed a slightly increased median survival (42 days, Figure 3A) compared to conventionally reared cohort (36 days, Figure S1A) axenic flies showed a greater difference in median survival between control and treated flies (39 days) than conventionally reared flies (30 days). There were also similar proportions of flies that had lost intestinal barrier function in axenic and conventional flies overexpressing PGRP-Lc at 20 and 30 days of age (∼10 and 50% respectively, Fig. S1B and Figure 3B).

**Figure 3:**
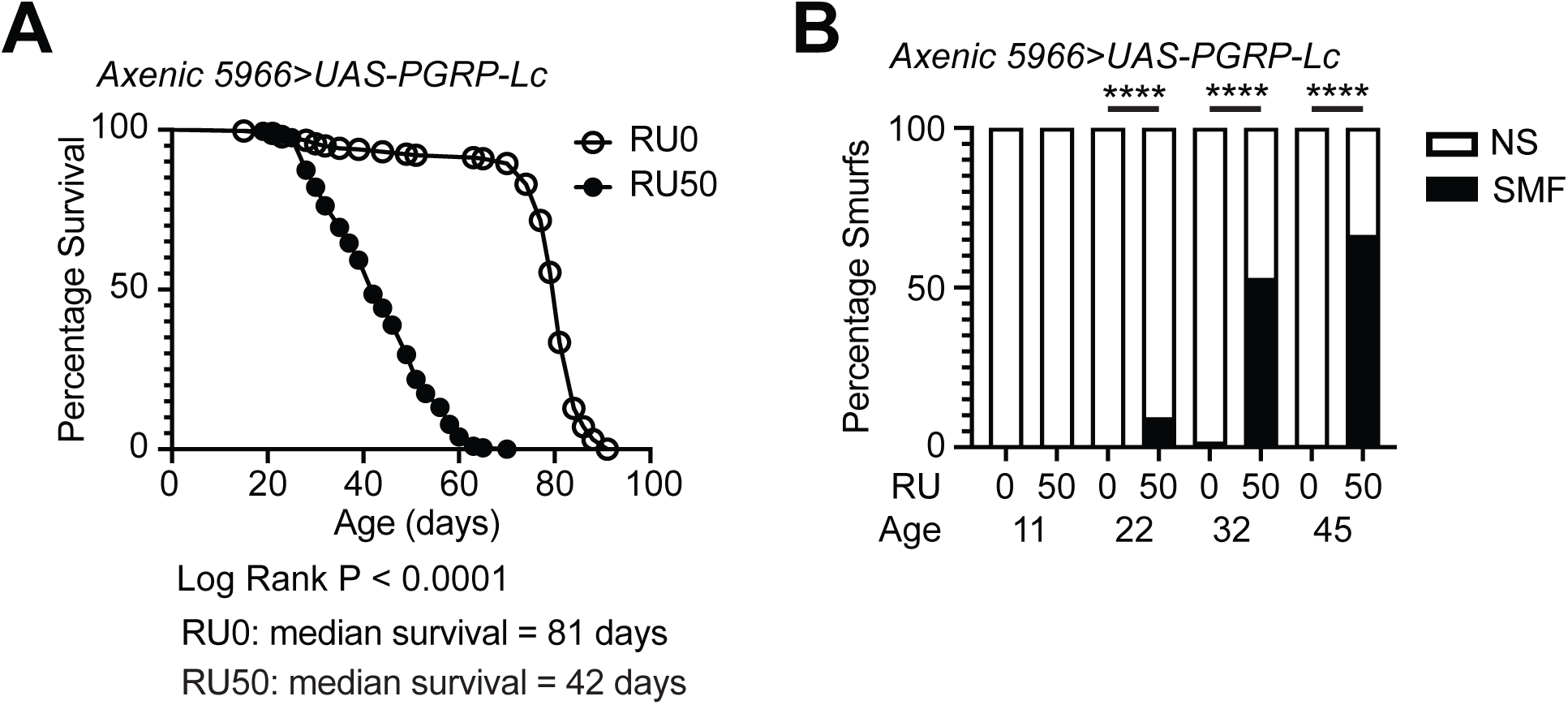
*PGRP-Lc* overexpression drives early mortality and intestinal barrier loss independent of the microbiota. (A and B) Lifespan curves (A) and Smurf proportions (B) for axenic *5966>UAS-PGRP-Lc* female flies drug induced (RU50) from early adulthood and uninduced controls (RU0). n > 200 flies/condition, NS = non-Smurf, SMF = Smurf. Log rank test was used for survival data and binomial test for Smurf proportions. *p < 0.05, **p < 0.01, ***p < 0.001, ****p < 0.0001.

To ensure that RU486 treatment was not itself driving these phenotypes we assessed mortality and intestinal barrier loss following RU486 treatment in 5966>w^1118^ flies. We found RU486 treatment had no consistent impact on survival or intestinal barrier loss in conventionally reared or axenic cohorts (Figure S2). These data demonstrate that *PGRP-Lc* overexpression rather than RU486 treatment drives mortality and intestinal barrier loss. We conclude from these data that the impact of *PGRP-Lc* overexpression on mortality and intestinal barrier loss is independent of the microbiota.

### Intestinal overexpression of PGRP-Lc impacts broad readouts of intestinal physiology

To assess the impact of PGRP-Lc overexpression on intestinal physiology we used The Ultimate Read of Dung (T.U.R.D.) (21) to profile the faecal deposits from non-Smurf females overexpressing PGRP-Lc compared to controls. We found *PGRP-Lc* overexpression had a significant impact on four measures of faecal output increasing the water content and alkalinity of deposits while reducing numbers of deposits (Figure 4A–E). This demonstrates that *PGRP-Lc* overexpression significantly impacts multiple elements of intestinal physiology prior to driving early intestinal barrier loss.

**Figure 4:**
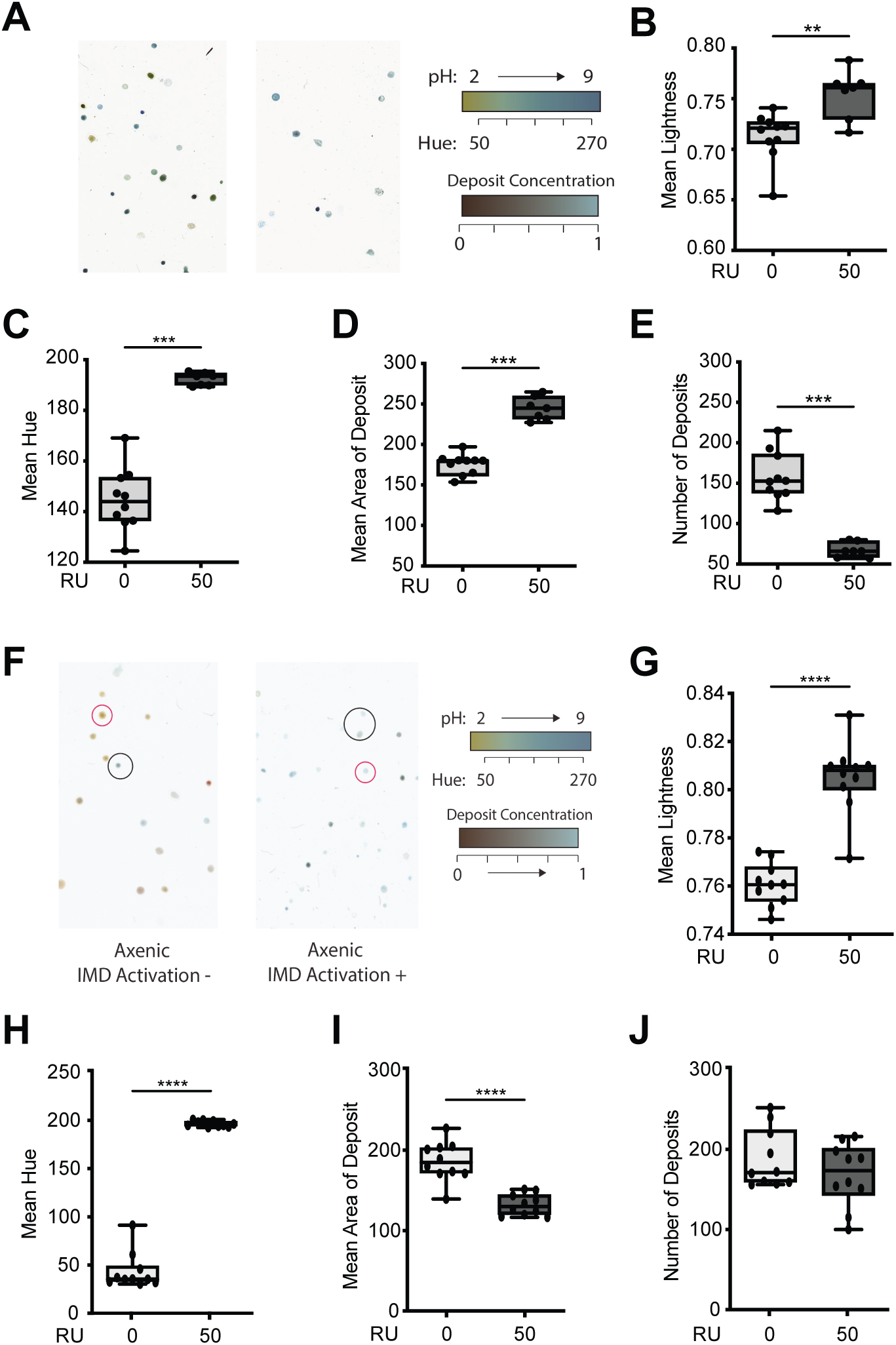
*PGRP-Lc* overexpression impacts intestinal physiology prior to barrier loss. (A – E) Representative T.U.R.D plate images (A) and analysis of faecal output (B-E) from conventionally reared *5966>UAS-PGRP-Lc* females at 30 days of age drug induced (RU50) from early adulthood and uninduced controls (RU0). (B) Mean lightness, (C) Mean hue, (D) Mean deposit area and (E) Number of deposits. (F – J) Representative T.U.R.D plate images (F) and analysis of fecal output (G-J) from axenic *5966>UAS-PGRP-Lc* females at 30 days of age drug induced (RU50) from early adulthood and uninduced controls (RU0). (G) Mean lightness, (H) Mean hue, (I) Mean deposit area and (J) Number of deposits. n = 10 plates with 10 flies/plate. Boxplots display the 25-75^th^ percentiles, with the horizontal bar at the median, and whiskers extending from the minimum to maximum points. Mann-Whitney U *p < 0.05, **p < 0.01, ***p < 0.001, ****p < 0.0001.

To assess whether microbial overgrowth in flies overexpressing PGRP-Lc drives these changes in intestinal physiology we repeated T.U.R.D. analysis on faecal matter from axenic individuals overexpressing PGRP-Lc compared to controls. We found that axenic flies overexpressing *PGRP-Lc* showed increases in faecal water content and alkalinity (Figure 4G & H) like conventional flies (Figure 4B & C). This suggests that the impact of *PGRP-Lc* overexpression on these measures is independent of the microbiota. However, in contrast to conventional flies, axenic flies showed a decrease in faecal deposit area and no significant change in deposit number (Figure 4I & J). This suggests that some aspects of the physiological changes that result from PGRP-Lc overexpression are driven by changes in the microbial population while others are independent.

### PGRP-Lc overexpression suppresses cell junction gene expression and drives stem cell proliferation

Previous work has shown intestinal barrier loss is accompanied by disruption in cell junction gene expression. Genetic knockdown of cell junction gene expression is sufficient to drive intestinal barrier loss, and the epithelial dysplasia associated with it (15,22–24). To assess the impact of intestinal immune activation on these phenotypes we first measured expression levels of the cell junction genes Snakeskin (SSK) and Mesh in *TiGS>UAS-PGRP-Lc* flies both before and after the onset of intestinal barrier loss. We found that *PGRP-Lc* overexpression reduced the expression levels of both SSK and Mesh in the intestines of 25-day old flies (Figure 5A & B). Importantly, we found suppression of SSK and Mesh expression in the intestines of flies that had not yet lost intestinal barrier function as well as those that had (Figure 5A & B). This demonstrates reduced expression of Snakeskin and Mesh occurs prior to the loss of intestinal barrier function in flies overexpressing *PGRP-Lc*.

**Figure 5:**
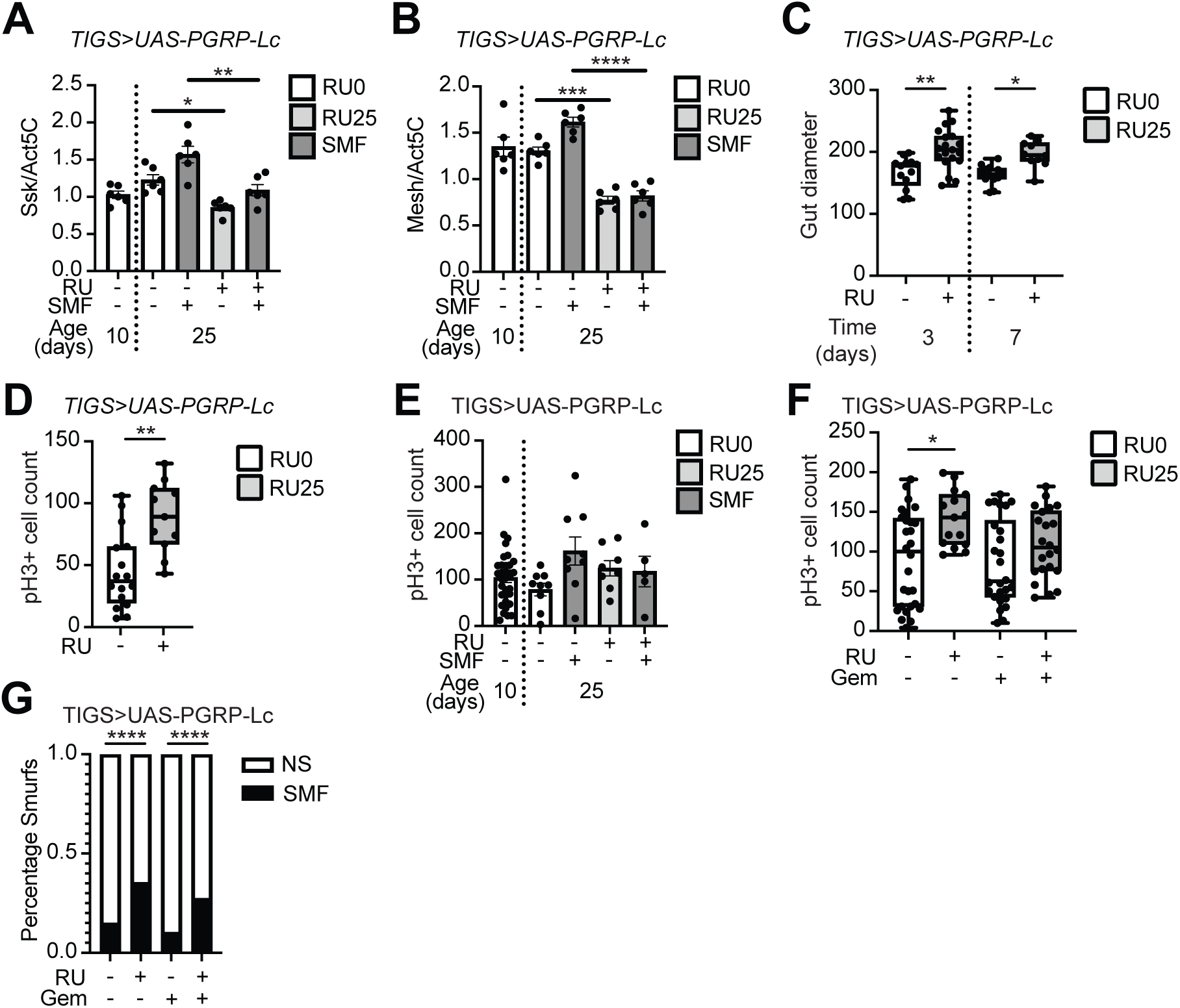
PGRP-Lc overexpression suppresses cell junction gene expression and drives stem cell proliferation. (A & B) Normalised mRNA level for Snakeskin (A, *Ssk)* and Mesh (B) in dissected whole gut samples from *TiGS>UAS-PGRP-Lc* non-Smurf and Smurf females fed RU486 and controls. n = 6 samples, 5 guts/sample. (C & D) Gut diameter at days 3 and 7 (C) and pH3+ cell counts at day 7 (D) post RU486 feeding from *TiGS>UAS-PGRP-Lc* non-Smurf females. (E) pH3+ cell counts at 25 days of age (15 days post RU486 feeding) from *TiGS>UAS-PGRP-Lc* non-Smurf and Smurf females fed RU486 and controls. N = (F & G) pH3+ cell counts in non-Smurf *TiGS>UAS-PGRP-Lc* female midguts at day 7 post RU486 and Gemcitabine feeding (F) and Smurf proportions at 15 days post RU486 and Gemcitabine feeding (23 days old) (G).Bar graphs show mean ± SEM. Boxplots display the 25-75^th^ percentiles, with the horizontal bar at the median, and whiskers extending from the minimum to maximum points. Two-way Anova with Tukey’s multiple comparisons (A, B & E). Mann Whitney U (C & D) Kruskal-Wallis with Dunn’s post hoc test (F) Binomial test (G) *p < 0.05, **p < 0.01, ***p < 0.001, ****p < 0.0001.

Induction of intestinal barrier loss through knockdown of Snakeskin has been shown to cause increased gut diameter and to drive stem cell proliferation (Salazar 2018). Therefore, we assessed the impact of *PGRP-Lc* overexpression on these phenotypes. We found an increased gut diameter in flies overexpressing *PGRP-Lc* compared to controls following just three days of treatment with RU486 (Figure 5C). Further, we found that *PGRP-Lc* overexpression resulted in increased cell proliferation, measured by increased phospho-Histone H3 positive (pH3+) cell numbers, in the intestine following 7 days of RU486 treatment (Figure 5D). No significant increase in cell proliferation was seen at later timepoints (Figure 5E), suggestive of stem cell exhaustion. Stem cell exhaustion has been reported in aged flies (25).

Given the close association between stem cell proliferation, the development of epithelial dysplasia, and the onset of intestinal barrier loss we wanted to assess whether stem cell over proliferation was required for intestinal barrier loss. To do this we fed *PGRP-Lc* overexpressing flies the chemotherapeutic drug Gemcitabine, which has previously been shown to limit cell proliferation in the Drosophila gut in a tumorigenesis model (26). Following 7 days of *PGRP-Lc* overexpression accompanied by Gemcitabine feeding we found a significant increase in cell proliferation only in *PGRP-Lc* overexpressing flies that were not fed Gemcitabine (Figure 5F); however, we note the variability in our data. We found no significant impact of gemcitabine feeding on the onset of intestinal barrier loss (Figure 5G). Given the minimal suppression of cell proliferation by gemcitabine feeding evident from our data we are unable to conclude that further suppression of cell proliferation would not impact the onset of intestinal barrier loss.

### Both JNK and NFkB signalling can drive intestinal barrier loss and mortality

Like mammalian Tumor Necrosis Factor (TNF), IMD signalling activates both its downstream NFκB transcription factor Relish and the adjacent Jun N-terminal Kinase (JNK) signalling pathway. The JNK pathway branches from the IMD pathway and contributes to cell proliferation and apoptosis in response to tissue damage caused by stress (27) Crosstalk between the two pathways occurs via a shared intermediary kinase (28), allowing for coordinated regulation of these outcomes.This means that the consequences of *PGRP-Lc* overexpression could be due to Relish activity, JNK signalling or both. To assess the impact of Relish and JNK activation on intestinal barrier loss and mortality we expressed constitutively active versions of *Relish* (*RelVP16*) and the Jun kinase kinase *hemipterous* (*hep^CA^*) using the TiGS driver. We confirmed activation by measuring expression levels of the target genes *Diptericin* (Relish) and Kay (JNK) (Figure S3). We found that both *TiGS>UAS-RelVP16* and *TiGS>UAS-hep^CA^* flies showed early mortality (Figure 6A & C) and intestinal barrier loss (Figure 6B & C). The impact of *UAS-hep^CA^* expression was particularly pronounced and led to rapid mortality and intestinal barrier loss like that reported for *Ssk* knockdown (15).

**Figure 6:**
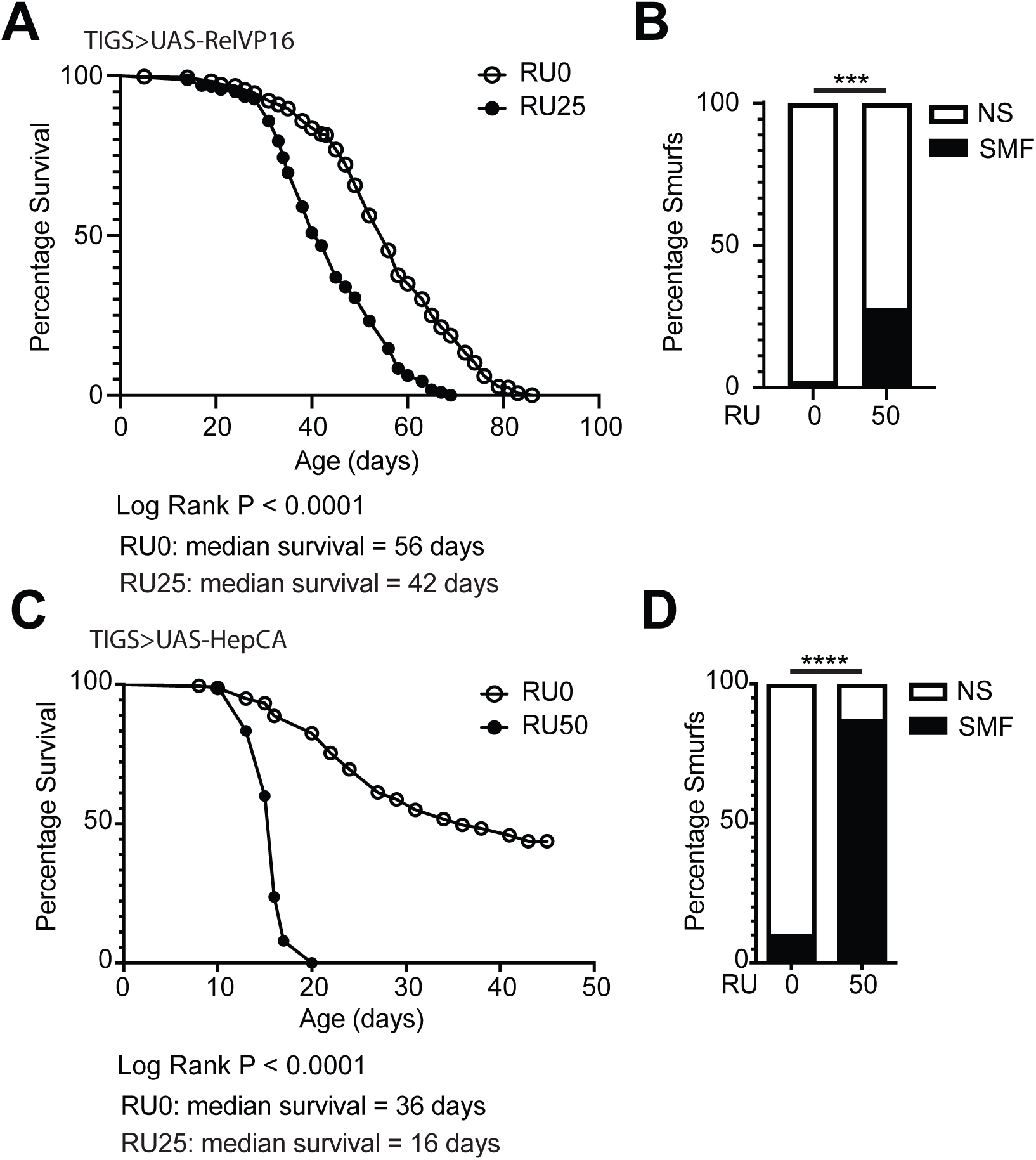
Both Relish and JNK activation drive early mortality and intestinal barrier loss. (A - D) Lifespan curves (A & C) and Smurf proportions (B & D) for *TiGS>UAS-RelVP16* (A & B) and *TiGS>UAS-hep^CA^* (C & D) female flies drug induced (RU25 or 50) from early adulthood and uninduced controls (RU0). n > 200 flies/condition, NS = non-Smurf, SMF = Smurf. Log rank test was used for survival data and binomial test for Smurf proportions. *p < 0.05, **p < 0.01, ***p < 0.001, ****p < 0.0001.

## Discussion

In this study we have clarified the relationship between innate immune signalling, intestinal barrier failure, microbial dysbiosis, epithelial dysplasia and mortality. Strikingly, by distinguishing the effects of immune activation before and after barrier failure we have shown that chronic immune activation has effects distinct from those caused by barrier loss but is nonetheless surprisingly well tolerated.

### Intestinal barrier failure and immune activation have distinct impacts on the microbiota

Previous studies in flies and mice have demonstrated that immune suppression can prevent age-related microbial dysbiosis (6,17,18). Conversely, immune activation has been shown to drive microbial dysbiosis (16,19). However, the contribution of intestinal permeability to microbial dysbiosis in this context has not been established. Interestingly, a previous study in *Drosophila* demonstrated that acute intestinal IMD activation for 7 days did not result in significant changes to the microbial population (29). This suggests that the alterations to the intestinal microbiota that are correlated with age-related immune activation may not be directly driven by immune activation but may rather result from decline of the intestinal epithelium.

Here we have addressed this by comparing microbial loads following IMD activation both before and after intestinal barrier failure. We find that flies carrying their native microbiota do show increased microbial loads following ten days of immune activation and prior to intestinal barrier loss. However, microbial loads show a further increase in flies with barrier failure. Interestingly, in axenic flies reassociated with Lactiplantibacillus plantarum and/or Acetobacter pasteurianus the microbial population does not respond to IMD activation and only shows overgrowth following intestinal barrier failure. Taken together with previous studies our data suggest that the impact of IMD activation on the microbial population is not acute and is species or strain specific. In contrast, intestinal barrier failure results in rapid microbial overgrowth that does not appear to be strain or species specific. This aligns with previous work demonstrating that while the microbial population shows some age-related changes prior to intestinal barrier loss, the largest changes, particularly to microbial load, occur after intestinal barrier failure (14). This supports the suggestion that much of the impact of intestinal inflammation on the microbial population may be through its impact on the intestinal barrier. Available data fits a model whereby inflammation drives initial changes to the microbial population that then exacerbate or prolong local inflammatory conditions ultimately causing barrier failure which results in a catastrophic loss of regulation of the microbial population. If this is the case it becomes critical to identify the microbial species that respond to inflammation in the absence of barrier loss if we are to identify effective targets for early intervention and health maintenance.

It is also important to identify why barrier failure renders the intestine unable to regulate it’s microbiota. In our model barrier failure doesn’t result in a worsening of immune activation but does drive further change to the microbial population. This means we must look beyond inflammation to identify the drivers of microbial dysbiosis in the inflamed intestine.

### Innate immune signalling drives barrier failure and death in the absence of the microbiota

Several studies have demonstrated that reducing immune activation (17,18) or removing the microbiota (6,14) improves both intestinal barrier function and survival. However, these studies have not addressed the relative impacts of immune activation or microbial dysbiosis on the intestinal barrier because manipulating one also impacts the other. Here we demonstrate that chronic immune activation results in intestinal barrier failure and death even in the absence of the microbiota. The role of immune activation in driving tissue damage and mortality is supported by recent work in a *Drosophila* tumour model that demonstrated that IMD activation resulting from bacterial translocation from the intestines of tumour hosts drove shortened lifespan (30). Genetic knockdown of the IMD receptor PGRP-Lc specifically in nephrocyte cells was sufficient to extend lifespan (30). This suggests that targeting immune activation directly, without removing the microbiota, may be sufficient to improve health outcomes following intestinal barrier loss.

We do not know whether acute immune activation results in small increases in intestinal permeability that cannot be detected by the Smurf assay. However, it’s important to note that measurable intestinal permeability does not arise immediately after immune activation. Instead, the onset of intestinal permeability retains its close association with the onset of mortality in our *Drosophila* model. This association between the Smurf phenotype and mortality has been well documented in *Drosophila* and other model systems (4,5,14,15). This delay in the onset of intestinal permeability suggests that this phenotype may result from ongoing epithelial damage.

### Immune activation drives intestinal stem cell proliferation prior to intestinal barrier failure

The mechanisms by which immune activation drives intestinal permeability remain unclear. One hypothesis is that this occurs through the impact of immune activation on intestinal stem cell (ISC) proliferation and subsequent epithelial dysplasia. Recent studies have demonstrated that loss of cell-cell contacts is sufficient to drive stem cell proliferation (15,22–24) and disrupt cell differentiation (23,24,31). In addition, it has been shown that septate junction components regulate Yorkie signalling and their loss enables Yorkie to drive expression of Unpaired 3 (Upd3) to promote ISC proliferation (32). IMD activation can also drive ISC proliferation and disrupt cell differentiation (29,33,34) and supressing IMD activation can reduce ISC proliferation (18,35,36).

Here, we further demonstrate that IMD activation induces ISC proliferation at an early timepoint, significantly preceding the onset of intestinal barrier loss. It is possible that this hyperproliferation leads to an accumulation of misdifferentiated cells, as is seen in the aged intestine (37). Cell junctions are dynamic as new daughter cells differentiate and integrate into the epithelium (38). Therefore, accumulation of misdifferentiated cells may impact cell junction formation and barrier function. Whether cell proliferation and subsequent dysplasia are necessary for intestinal barrier failure remains an important question.

In addition to over proliferation of ISCs and disrupted cell differentiation, aberrant enterocyte shedding may also be a potential driver of epithelial damage that could cause barrier loss. It has been shown that infection induced enterocyte shedding requires IMD and activation of IMD signalling is sufficient to induce shedding (39). Enterocyte shedding was also accompanied by increased ISC proliferation (39), however, the impact of this rapid epithelial turnover on cell-cell contacts and barrier function is unknown.

### Both NFκB and JNK signalling can drive intestinal barrier failure and mortality

It has been well established that JNK activity drives age-related ISC proliferation in *Drosophila* (37,40) and is important for microbial induction of proliferation (41). Inhibition of JNK has been shown to reduce ISC proliferation following genetic induction of intestinal barrier failure (24) or dextran sodium sulfate (DSS) induced inflammation (42). Following DSS treatment and in a tumorigenesis model inhibition of JNK also improved intestinal barrier function (42,43). JNK has also been shown to regulate intestinal barrier function in *C. elegans* (44). Further, both JNK and IMD signalling can induce enterocyte cell shedding and inhibition of JNK reduces cell shedding following infection (39). That JNK signalling regulates both ISC turnover and enterocyte shedding likely explains the rapid onset of intestinal barrier failure and death when JNK is constitutively activated.

The role of JNK activity following activation of IMD signalling is complicated by the fact that Relish is a negative regulator of JNK (45,46). This may explain the difference in response to constitutively active IMD versus JNK, whereby IMD activation appears well tolerated for some time before the onset of intestinal barrier failure and mortality. There are studies that have distinguished the roles of JNK and Relish activity following constitutive activation of IMD. For example, remodelling of *Drosophila* airways following IMD activation was dependent on JNK not Relish (47). Further, ISC proliferation induced by *PGRP-Lc* overexpression has previously been shown to be dependent on Relish (39).

There is also evidence that JNK and Relish (NFκB) may work cooperatively in some contexts, e.g. enterocyte cell shedding (39). How both JNK and Relish contribute to age-related intestinal permeability warrants further investigation.

### Conclusions

Our work demonstrates that animals are surprisingly resilient to chronic immune activation, in a way that they are not to the acute induction of intestinal barrier failure. Immune-induced intestinal barrier failure is more closely associated with the onset of mortality than with the onset of immune activation. Further, we add to building evidence that intestinal permeability is a key driver of microbial dysbiosis through mechanisms that are independent of and distinct from its impact on inflammation.

Preventing excessive inflammation is critical for health maintenance and has the potential to prevent or delay age onset intestinal permeability, as well as to address the consequences of a leaky gut. However, the impact of intestinal permeability on health needs to be considered independently of its relationship with inflammation.

## Materials and Methods

### Fly stocks

This work used the TiGS-2-GeneSwitch (GS) (48), 5966-GS (49), Daughterless-GS (DaGS), S106-GS (BDSC #8151) driver lines, UAS-RelVP16 (Bloomington Drosophila Stock Center (BDSC) #36547), UAS-Hep^CA^ (BDSC #9306, gift of Marc Dionne), UAS-PGRP-Lc (BDSC #30919) and w^1118^.

### Fly culture

Flies were maintained in either vials or bottles in incubators with 65% humidity and a 12-hour light/dark cycle, at either 18°C or 25°C. Stocks were maintained on cornmeal medium comprised of 1 % agar (SLS, FLY1020), 3% dried inactivated yeast (Genesee Scientific, 62-106), 1.9 % sucrose (Duchefa Biochemie, S0809), 3.8 % dextrose (Melford, G32040) and 9.1 % cornmeal (SLS, FLY1110), all given in wt/vol, and water at 90% of the final volume. Preservatives; 10 % Nipagin (SLS, FLY1136) in ethanol and an acid mixture comprised of 4.15 % phosphoric acid (VWR, 20624.262) and 41.8 % propionic acid (Sigma, P5561) in water were added following boiling.

### Lifespan

All experiments were carried out using age-matched, mated females, housed at a density of 27-32 flies per vial. These flies were flipped onto new food within 24 hours of eclosing. They were then allowed a further 24-48 hours to mate before the mated females were sorted into vials under light CO_2_ anaesthesia. Flies were flipped to fresh food every 2-3 days and the number of dead flies was counted. At least 8 vials of 30 females were used per lifespan. To control for bacterial growth on the food, control and experimental flies are always transferred to fresh food at the same time points.

Mifepristone or RU-486 (Cayman chemicals, 10006317), hereafter referred to as RU, food was prepared by the addition of 125 µL of 20 mg/ml RU stock solution per 100mL food for a final concentration of 25 µg/ml (RU25). To ensure sufficient volume for effective mixing 125 µL of RU stock was added to 125 µL of 100% ethanol per 100 mL of food prior to addition to the food. For a concentration of 50 µg/ml (RU50) 250 µL of RU stock solution was added per 100ml food. Control food was prepared by the addition of 250 µL ethanol per 100 mL food. Food was mixed thoroughly to ensure a homogenous solution.

Gemcitabine (Merck, APO456787943) was prepared at a concentration of 100 µM in water. 50 µl of this solution was added to the surface of standard cornmeal food and allowed to dry for 24 hours in the dark before use.

### Smurf assay

The smurf assay (Rera 2011) for intestinal barrier integrity was performed by flipping flies onto food containing 2.5 % FD&C blue food dye #1 for 24 hours (FastColours, 08BLU00201). After 24 hours, the flies were flipped back onto dye-free food and the number of ‘smurf’ and ‘non-smurf’ flies were recorded.

### Generation of axenic and reassociated flies

To generate sterile (axenic) flies, eggs were collected on apple juice agar plates seeded with a paste of live baker’s yeast in ddH_2_O, then washed and treated with bleach and ethanol as described previously (Bakula 1969). All steps were carried out in a laminar flow hood. Briefly, <14 hour-old embryos were dechorionated in 2 % sodium hypochlorite solution for 2.5 minutes, rinsed in 70 % ethanol for 5 minutes and then rinsed with ∼200 mL sterile PBS. Embryos were then resuspended in sterile 0.01 % PBSTx (PBS + 0.01 % Triton X-100 (Sigma, X100)) and transferred to sterile autoclaved cornmeal medium.

For re-association of sterilised embryos with specific bacteria, we obtained working bacterial cultures by inoculating Lactiplantibacillus plantarum (strain and source) in MRS media (supplier etc) and Acetobacter pastiurianus (strain and source) in Mannitol media (supplier or recipe etc) and culturing at 30°C for 48 hours. We first established the relationship between optical density at OD600 and bacterial colony forming units (CFU) per volume, to allow us to standardise CFU/fly vial. Bacterial cultures were centrifuged at 2000rpm for 5 minutes, supernatant was removed, and the bacterial pellet was resuspended in sterile 1xPBS. The optical density was adjusted to an OD600 of 0.2. The bacterial solution was then serially diluted 1:5 and plated on the respective bacterial agar. Following incubation for 24-48 hours, colonies were counted from a single dilution sample and this dilution factor was used to calculate CFU/mL. 24-hours prior to embryo collection bacterial cultures prepared as described above and adjusted to the appropriate dilution were used to evenly inoculate vials containing 8ml of sterile cornmeal medium with 3×10^6^ CFU/vial. Vials inoculated with bacteria were then seeded with sterile embryos. To maintain the integrity of the bacterial population introduced, adult reassociated progeny were handled as though axenic All medium for sterile and reassociated flies was sterilised by autoclaving for 15 minutes at 121 °C, and all equipment used to push flies was first sterilised in Virkon for at least 20 minutes, then rinsed and stored in 80 % ethanol. Gas pads were subsequently dried by running CO_2_ through them. To maintain sterility or reassociation, females were separated into lifespan vials next to an open flame and flipped to new sterile food every 2-3 days in a laminar flow hood.

### Bacterial DNA extraction

30 whole flies were sorted under light CO_2_ anaesthesia into microcentrifuge tubes and frozen at −80 °C. Flies were then surface sterilised in 70 % ethanol for 5 minutes and rinsed twice in sterile PBS. 5 flies were sorted into sterile microcentrifuge tubes and homogenised in 200 µl lysis buffer (1 X TE, 1 % Triton X-100, 1/100 proteinase K (NEB, P8107S)) using a mechanical pestle, with a further 300 µl of lysis buffer being added after homogenisation. Lysate was incubated for 1.5 hours at 55 °C then transferred to screw cap vials containing 0.4 g sterilised low binding 100-micron silica beads (OPS Diagnostics, BLBG 100-200-11) and vortexed for 10 minutes at max speed. Lysate was then transferred back to the original tube and incubated for a further 1.5 hours at 55 °C followed by 10 minutes at 95 °C.

### RNA extraction

Total RNA was extracted from 5 midguts per biological replicate. Midguts were dissected in ice-cold PBS and included the cardia and midgut, through to the midgut-hindgut junction, excluding the hindgut and Malpighian tubules. Midguts were homogenised in 100 µL TRIzol™ (Thermofisher, 15596026) using a motorised pestle. RNA was extracted following the standard TRIzol protocol.

### cDNA preparation

For a 10 µL reaction, 1 µL DNase buffer (Thermo, EN0521) and 0.5 µL DNase 1 (Thermo, EN0521) were added to 8.5 µL RNA solution and incubated for 30 minutes at 37 °C, to remove any DNA contaminants. The DNase enzyme was then deactivated by the addition of 0.5 µL 50mM EDTA (Thermo, EN0521) and subsequent incubation for 10 minutes at 65 °C. cDNA synthesis was carried out using Thermo Scientific’s First Strand cDNA Synthesis kit. In short, 1 µL of random hexamers (Thermo, SO142) were added and the samples were incubated at 70 °C for 5 minutes before a 9 µL of a master mix containing 4 µL first strand buffer (Thermo, EPO442), 0.5 µL RIbolock RNase inhibitor (Thermo, EO0382), 2 µL 10mM dNTPs (Thermo, R0181), 0.2 µL RevertAid reverse transcriptase (Thermo, EPO442) and 2.3 µL molecular grade water (Sigma, W4502) per well, was added. The samples were then incubated for 10 minutes at room temperature, followed by 1 hour at 37 °C and 10 minutes at 70 °C. The samples were spun following each incubation step.

### Quantitative PCR

qPCR was performed with Power SYBR™ Green master mix (Fisher, 10219284) on a Bio-RAD CFX Connect Real-Time PCR detection system. Either 4.5 µL bacterial DNA or 1 µL of *Drosophila* cDNA sample was used in a 10 µL reaction with 5 µL of Power SYBR™ Green and 0.5 µL of L + R primer mix. Any remaining volume was made up with molecular grade water.

Cycling conditions were: 95°C for 10 minutes, then 40 cycles of 95°C for 15 seconds followed by 60°C for 60 seconds. All calculated gene expression values were normalised to the value of a housekeeping gene, either Act5C or rp49.

The primer sequences used to assess gene expression are as follows:

SSK_F – CACTGGATGCCACACCATT

SSK_R – TGGTGTCGCACAGCTCTC (15)

Mesh_F – AGCCCGATCAATACTCAGGA

Mesh_R – CCATATACCAGGCCAGAGGA (15)

Dpt_F – ACCGCAGTACCCACTCAATC

Dpt_R – CCCAAGTGCTGTCCATATCC

PGRP_49F – CCTTCCTGCTGGGTATCGTA

PGRP_49R – CCACCAATGTTGTCCAGATTT

Kay_F – CAGCATCAGCGACAGGATTA

Kay_R – TCTGGCCGGTCTCAAAGTT (43)

Act5c_F – TTGTCTGGGCAAGAGGATCAG

Act5c_R - ACCACTCGCACTTGCACTTTC

p49_F – ATCGGTTACGGATCGAACAA

Rp49_R - GACAATCTCCTTGCGCTTCT

Universal primers for the 16S ribosomal RNA gene were against variable regions 1 (V1F) and 2 (V2R) (Claesson 2010). Levels of the bacterial 16S rRNA gene were normalised to the level of the *Drosophila* Actin gene, as previously published (Clark 2015).

16S_V1F - AGAGTTTGATCCTGGCTCAG

16S_V2R – CTGCTGCCTYCCGTA

Act_F – TTGTCTGGGCAAGAGGATCAG

Act_R - ACCACTCGCACTTGCACTTTC

### Faecal analysis

Faecal analysis was conducted as described previously (14).

### Dissection and immunofluorescent staining of midguts

Whole guts were dissected into ice-cold PBS, with the crop, cardia, ovaries, and Malphigian tubules removed. Fixation of was carried out with 16% PFA for 30 minutes, followed by three immediate washes in PBS. Blocking buffer (PBS with 0.5% Triton X-100, 0.5% BSA and 5% fresh NGS) was added and incubated for 30 minutes.

Samples were immunostained with primary antibody (rabbit a-pH3, Merck, H0412) at 1:2000 overnight on a rocker at 4°C. The following morning, guts were washed three times for 10 minutes each, in PBS. Samples were incubated with the secondary antibody (Goat anti-rabbit 555, supplier) at 1:500 on a rocker at room temperature for 2 hours. DAPI (1:2000) was added for a 10-minute incubation to stain nuclei. Guts were again washed three times for 10 minutes each, in PBS. Samples were mounted in VectaShield Antifade Mounting Medium (Vector Laboratories, H-1000-10)

### Microscopy

For the quantification of pH3+ cells, images were acquired with a 10x objective and rhodamine laser under the Zeiss Cell Observer microscope using AxioVision 40 software (Carl Zeiss). The total number of pH3+ cells across the entire length of each gut was counted.

To measure variations in midgut diameter, single images were taken under the 10x objective and DAPI laser using the Zeiss Cell Apotome microscope and measured using the AxioVision Rel 4.8 software (Carl Zeiss). Diameter was calculated just above the midgut-hindgut junction for each midgut, as in (15).

### Statistics

All Statistical analyses were performed using GraphPad Prism 9, excluding the binomial tests which were performed in R (Rstudio build 372, R version 4. 0. 3). Survival curve analysis was completed using the log-rank test in GraphPad Prism; the median survival was calculated by Graph Pad Prism 8 and is an approximation to the closest sampling time. Normality of the data was tested by Shapiro-Wilk tests to determine the most appropriate statistical test. Differences between two points were analysed by students t-test, or Mann-Whitney tests for normally and non-normally distributed data respectively. Differences in gene expression values between more than two time points, within or between conditions, were analysed by two-way Anova with Tukeys multiple comparison or Kruskal-Wallis with Dunn’s test for normally and non-normally distributed data respectively. P-values less than 0.05 were considered statistically significant. Bar-graph error bars depict ± standard error of the mean (SEM). For all box-plots shown, boxes display the 25-75^th^ percentiles, with the horizontal bar at the median, and whiskers extending from the minimum to maximum points.

## Acknowledgements

We thank Marc Dionne and the Bloomington Drosophila stock center for fly stocks and H. Siddle for technical support. This work was supported by the Durham Doctoral Scholarship program. Stocks obtained from the Bloomington Drosophila Stock Center (NIH P40OD018537) were used in this study.

## Author Contributions

Conceptualisation, R.I.C. and D.W.W.; Methodology, J.A., C.K, C.H. and R.I.C.; Investigation, J.A., C.K, C.H. and R.I.C.; Formal Analysis, J.A., C.K, C.H. and R.I.C.; Writing – Original Draft, J.A. and R.I.C.; Writing – Review and Editing, J.A., D.W.W., R.I.C and D.P.D.; Supervision, R.I.C and D.P.D.

---

**Figure S1:**
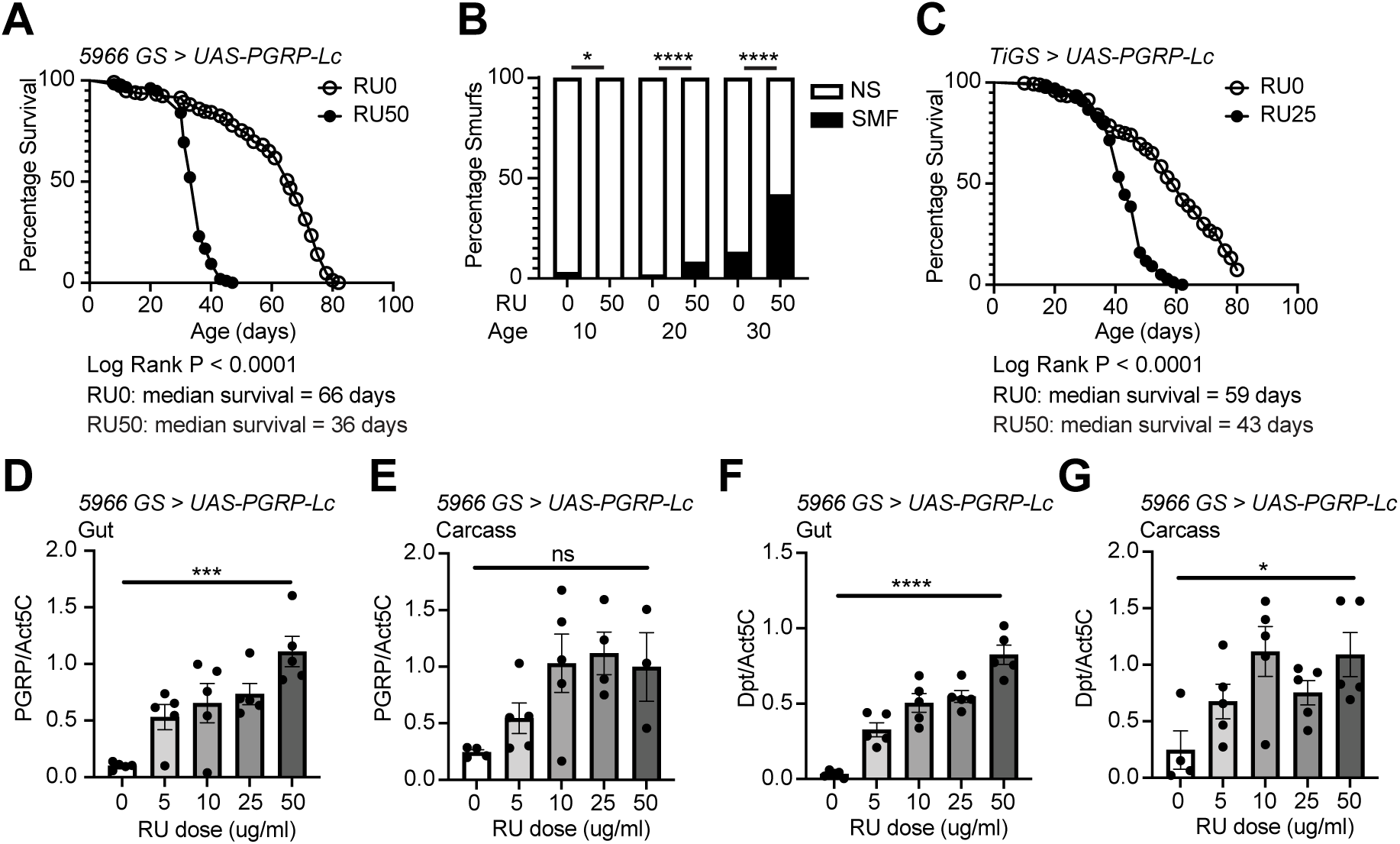
The 5966 geneswitch drives UAS-PGRP-Lc expression in the gut and other tissues which results in early mortality and intestinal barrier loss. (A and B) Lifespan curves (A) and Smurf proportions (B) for *5966>UAS-PGRP-Lc* female flies drug induced (RU50) from early adulthood and uninduced controls (RU0). n > 200 flies/condition, NS = non-Smurf, SMF = Smurf. (C) Replicate lifespan curve for *TIGS>UAS-PGRP-Lc* female flies drug induced (RU25) from early adulthood and uninduced controls (RU0). n > 200 flies/condition. Log rank test was used for survival data and binomial test for Smurf proportions. (D-G) Normalised mRNA level for *PGRP-Lc* (D and E) and *diptericin (Dpt*, F and G)) in dissected whole gut (D and F) and body carcass samples (E and G) on diXerent RU486 doses. n = 6 samples, 5 guts/sample and 5 carcasses (gut removed)/sample. Bar graphs show mean ± SEM. Two-way Anova with Tukey’s multiple comparisons. *p < 0.05, **p < 0.01, ***p < 0.001, ****p < 0.0001.

**Figure S2:**
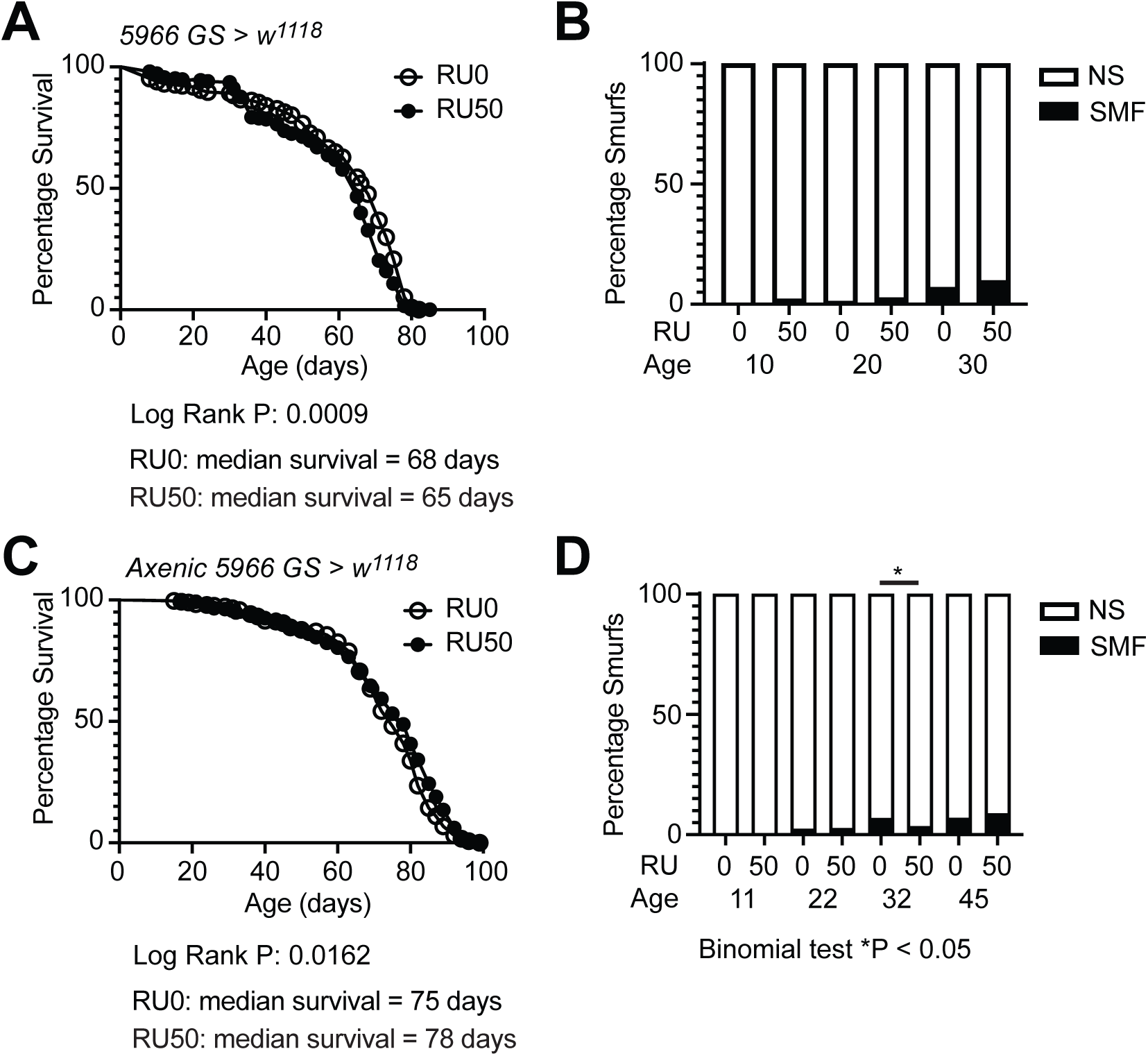
RU486 feeding doesn’t consistently drive early mortality or intestinal barrier loss. Lifespan curves (A and C) and Smurf proportions (B and D) for conventional (A and B) axenic (C and D) *5966>w^1118^* female flies drug fed (RU50) from early adulthood and controls (RU0). n > 200 flies/condition, NS = non-Smurf, SMF = Smurf. Log rank test was used for survival data and binomial test for Smurf proportions. *p < 0.05, **p < 0.01, ***p < 0.001, ****p < 0.0001.

**Figure S3:**
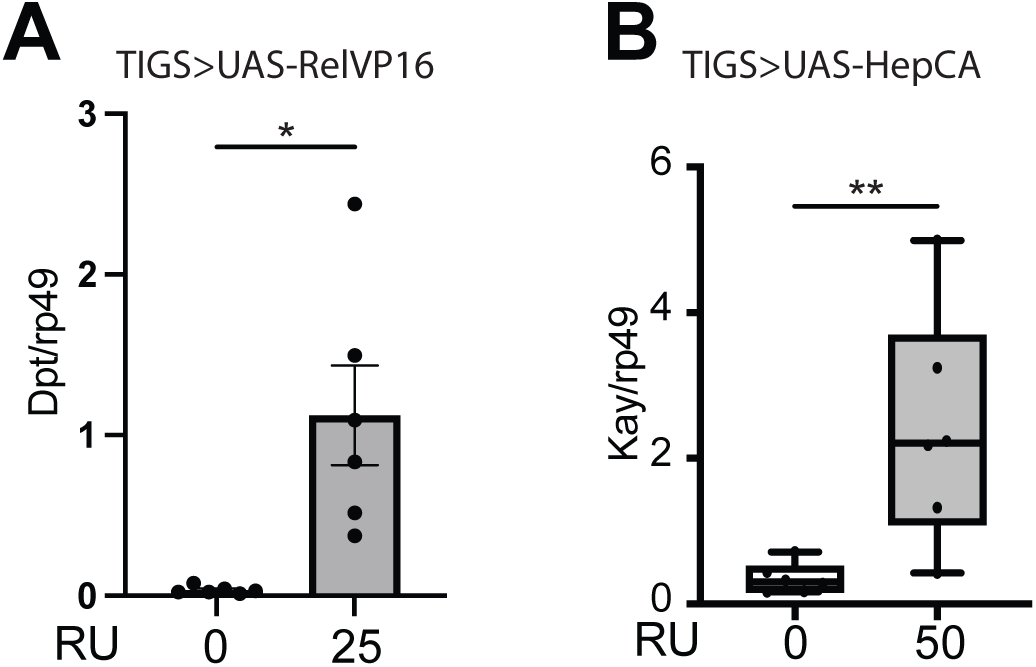
Constitutively active RelVP16 and HepCA drive target gene expression. (A & B) Normalised mRNA level for *Diptericin (Dpt)* (A) and *Kay* (B) in dissected whole gut from *TiGs>UAS-RelVP16* (A) and *TiGs>UAS-HepCA* (B) females fed RU486 for 72 hours or controls. n = 6 samples, 5 guts/sample. Bar graphs show mean ± SEM. Boxplots display the 25-75^th^ percentiles, with the horizontal bar at the median, and whiskers extending from the minimum to maximum points. Two-way Anova with Tukey’s multiple comparisons. *p < 0.05, **p < 0.01, ***p < 0.001, ****p < 0.0001.

